# Community-based benchmarking improves spike rate inference from two-photon calcium imaging data

**DOI:** 10.1101/177956

**Authors:** Philipp Berens, Jeremy Freeman, Thomas Deneux, Nicolay Chenkov, Thomas McColgan, Artur Speiser, Jakob H. Macke, Srinivas C. Turaga, Patrick Mineault, Peter Rupprecht, Stephan Gerhard, Rainer W. Friedrich, Johannes Friedrich, Liam Paninski, Marius Pachitariu, Kenneth D. Harris, Ben Bolte, Timothy A. Machado, Dario Ringach, Jasmine Stone, Luke E. Rogerson, Nicolas J. Sofroniew, Jacob Reimer, Emmanouil Froudarakis, Thomas Euler, Miroslav Román Rosón, Lucas Theis, Andreas S. Tolias, Matthias Bethge

## Abstract

In recent years, two-photon calcium imaging has become a standard tool to probe the function of neural circuits and to study computations in neuronal populations^1,2^. However, the acquired signal is only an indirect measurement of neural activity due to the comparatively slow dynamics of fluorescent calcium indicators^3^. Different algorithms for estimating spike rates from noisy calcium measurements have been proposed in the past^4–8^, but it is an open question how far performance can be improved. Here, we report the results of the *spikefinder* challenge, launched to catalyze the development of new spike rate inference algorithms through crowd-sourcing. We present ten of the submitted algorithms which show improved performance compared to previously evaluated methods. Interestingly, the top-performing algorithms are based on a wide range of principles from deep neural networks to generative models, yet provide highly correlated estimates of the neural activity. The competition shows that benchmark challenges can drive algorithmic developments in neuroscience.

## Introduction

Two-photon calcium imaging has become a standard tool to probe the function of neural circuits and to study computations in neuronal populations^1,2^. Indeed, the latest advances in scanning technologies make it now possible to record neural activity from hundreds or even thousands of cells simultaneously^9-11^. However, the resulting fluorescence signal is only an indirect measurement of the underlying spiking activity, as it reflects the comparatively slow cellular dynamics of cellular calcium and the fluorescent calcium indicators^3,4,12^. Thus, to relate large-scale population recordings to the spiking activity of neural circuits we fundamentally require techniques to infer spike rates from the fluorescent traces.

Over the past decade, a number of algorithms for solving this problem have been proposed. Many of them assume a forward generative model of the calcium signal and attempt to invert it to infer spike rates. Examples of this approach include deconvolution techniques^13,14^, template-matching^10,15^and approximate Bayesian inference ^4^’^7^’^8^. Such forward models incorporate a priori assumptions about how the measured signal is generated, e.g. about the shape of the calcium fluorescence signal induced by a single spike and the statistics of the noise. In contrast, comparatively few groups have attempted to solve the problem through supervised learning^6,16^, where a machine learning algorithm is trained to infer the spike rate from calcium signal using simultaneously recorded spike and calcium data for training.

Despite this progress, it is still an open question whether current algorithms already achieve the best possible performance for the task, or whether the observed performance can still be improved upon by algorithmic developments. To answer this question, we organized the *spikefinder* challenge. This challenge aimed at two goals: it was supposed to (1) provide a standardized framework to evaluate existing spike inference algorithms on identical data and (2) catalyze the development of new spike inference algorithms through crowd-sourcing. Such challenges have been used successfully in machine learning, computer vision or physics to drive algorithmic developments^17,18^. We present ten of the submitted algorithms which show improved performance compared to previously evaluated methods ^6^. Interestingly, the top-performing algorithms are based on a range of principles from deep neural networks to generative models, yet provide highly correlated estimates of the neural activity.

## Results

For the *spikefinder* challenge, we used five benchmark data sets consisting in total of 92 recordings from 73 neurons, acquired in the primary visual cortex and the retina of mice (see Table 1). In brief, data sets I, II and IV were collected with OGB-1 as a calcium dye, while data sets III and V were collected with the genetically encoded indicator GCamp6s. Similarly, there were differences in scanning method and scan rate between the data sets: For example, data set I was recorded using 3D AOD scanners at very high scan rates^9^, while data set II was recorded using conventional galvo-scanners at fairly low speed. For all data sets, calcium imaging had been performed simultaneously with electrophysiological recordings allowing to evaluate the performance of spike rate inference algorithms on ground truth data^6^. Importantly, all data was acquired at a zoom factor typically used during population imaging experiments, ensuring that all benchmark results reflect performance under the typical use-case conditions.

**Table 1.** Overview over datasets with training and test data used in the competition.

| Dataset | Scan method | Indicator | Avg. scan rate (Hz) | N in training set | N in test set |
| --- | --- | --- | --- | --- | --- |
| I | 3D AOD | OGB-1 | 322.5 | 11 | 5 |
| II | galvo | OGB-1 | 11.8 | 21 | 10 |
| III | resonant | GCamp6s | 59.1 | 13 | 6 |
| IV | galvo | OGB-1 | 7.8 | 6 | 3 |
| V | resonant | GCamp6s | 59.1 | 9 | 8 |

For the challenge, we split the data into a training and a test set, making sure that all recordings from a single neuron were either assigned to the training or the test set. For the training data, we made both the calcium and the spike traces publicly available, but kept the spike traces secret for the test data. Additionally, the publicly available data sets provided by the GENIE project^19^ were available as training data. This allowed participants to adjust their models on the training data set, while avoiding overfitting to the specific benchmark data set providing a realistic estimate of the generalization performance. Participants could upload predictions for the spike rate generated by their algorithm on a dedicated website (see Methods) and see their performance on the training set during the competition phase. Results on the test set were not accessible to the participants during the competition. The primary evaluation measure for the competition was the Pearson correlation coefficient between the true spike trace and the prediction sampled at 25 Hz (equivalent to 40 ms time bins) as previously described^6^.

We obtained 37 submissions, from which we selected all algorithms performing better than the spike-triggered-mixture model algorithm (STM), which had previously been shown to outperform other published algorithms on this data^6^. In addition, if there were multiple submissions from the same group, we used the one with the highest correlation on the test set. This resulted in a total of ten algorithms that we studied in greater detail and that are included in this paper^1^ (see Table 2).While seven of these algorithms were designed specifically for the purpose of the challenge, three were heavily based on methods published previously (see Table 2 for overview).

**Table 2.**
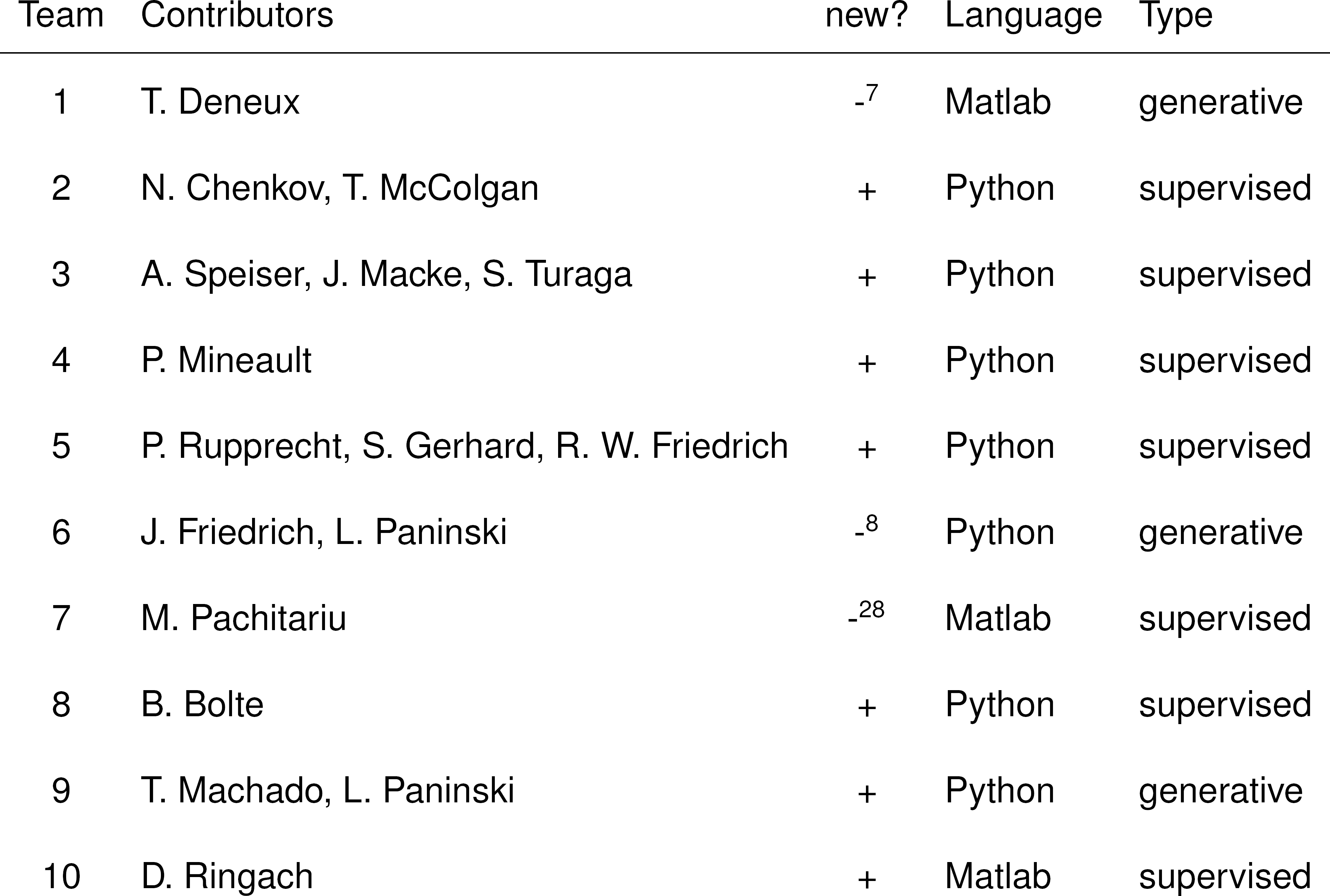
Overview over submitted algorithms and key properties.

Interestingly, these submissions include algorithms based on very different principles: some of the algorithms built on the classical generative models of spike-induced calcium dynamics^4^, while others relied on purely data-driven training of deep neural networks or pursued hybrid strategies (see Table 2). Algorithms based on generative models of the calcium fluorescence include the MLspike algorithm by Team 1^7^, which performs efficient Bayesian inference in a biophysical model of measured fluorescence including a drifting baseline and nonlinear calcium to fluorescence conversion (for a detailed description of each algorithm, see Appendix). Within the same group of algorithms, Team 6 took a decidedly different approach, approximating the calcium fluorescence by an autoregressive process and finding the spike trains by solving a non-negative sparse optimization problem^8,20^. A similar approach is taken by Team 7, who use L_0_-deconvolution in a linear model of calcium fluorescence with exponential calcium filters.

In contrast, many other algorithms took a purely data-driven approach^6^ and trained different variants of deep neural networks to learn the relationship between measured spike and calcium traces. For example, the algorithm by Team 2 used a straightforward network architecture with eight convolutional layers with consecutively smaller convolutional filters and one intermediate recurrent LSTM layer. The filters learned in the first layer provide a rich basis set for different spike-calcium relationships (see Fig. 5). Similarly, the algorithm by Team 5 used fairly standard components, consisting of convolutional and max-pooling layers. In contrast, the algorithms proposed by Teams 3, 4, and 8 combined more involved elements such as residual blocks^21^ or inception cells^22^. The key features of the different DNN-based approaches are summarized in Table 3.

**Table 3.** Overview over different strategies used by DNN-based algorithms. Architecture briefly summarizes main components. conv: convolutional layers, typically with nonlinearity; lstm: recurrent long-short-term memory unit; residual: residual blocks; max: max-pooling layers; inception: inception cells. For details, refer to the descriptions of the algorithms in the supplementary material.

| Team | Architecture | Optimizer | Dropout | Cost | dataset specific |
| --- | --- | --- | --- | --- | --- |
| 2 | conv / lstm | Adam | yes | correlation | indicator |
| 3 | RNN/CNN | Adam |  | cross-entropy | separate |
| 4 | residual / lstm | Adam | yes | scaled SSE | transfer |
| 5 | conv / max | Adagrad | no | MSE | embedding |
| 8 | inception | Adam | yes | correlation | embedding |

The best algorithm increased the average correlation on the test set from 0.36 by 0.08 to 0.44 compared to the STM (Figure 1A, B; Table 4). This corresponds to an increase of more than 40% in variance explained for the best algorithms, similar to the improvement seen between the STM algorithm and f-oopsi (see Table 4 and ref. ^6^). For all algorithms, performance varied substantially between data sets with the best results observed on data set I. Interestingly, performance gains were typically larger on GCaMP6 than on OGB-1 data sets (Figure 1B). Surprisingly, the top group of six algorithms performed equally well, despite using very different methodologies. Indeed, when we computed a repeated measures ANOVA, we were not able to distinguish the first six algorithms during post-hoc testing (Figure 1C). In addition, we evaluated to what extent the algorithms overfitted the training data. For example, it is possible that algorithms extracted peculiarities of the training data that did not transfer to the test data, resulting in artificially high correlation coefficients on the training data. We found that most algorithms showed similar performance for both the training and the test set, with evidence for overfitting in some of the DNN-based algorithms (Figure 1D).

**Figure 1:**
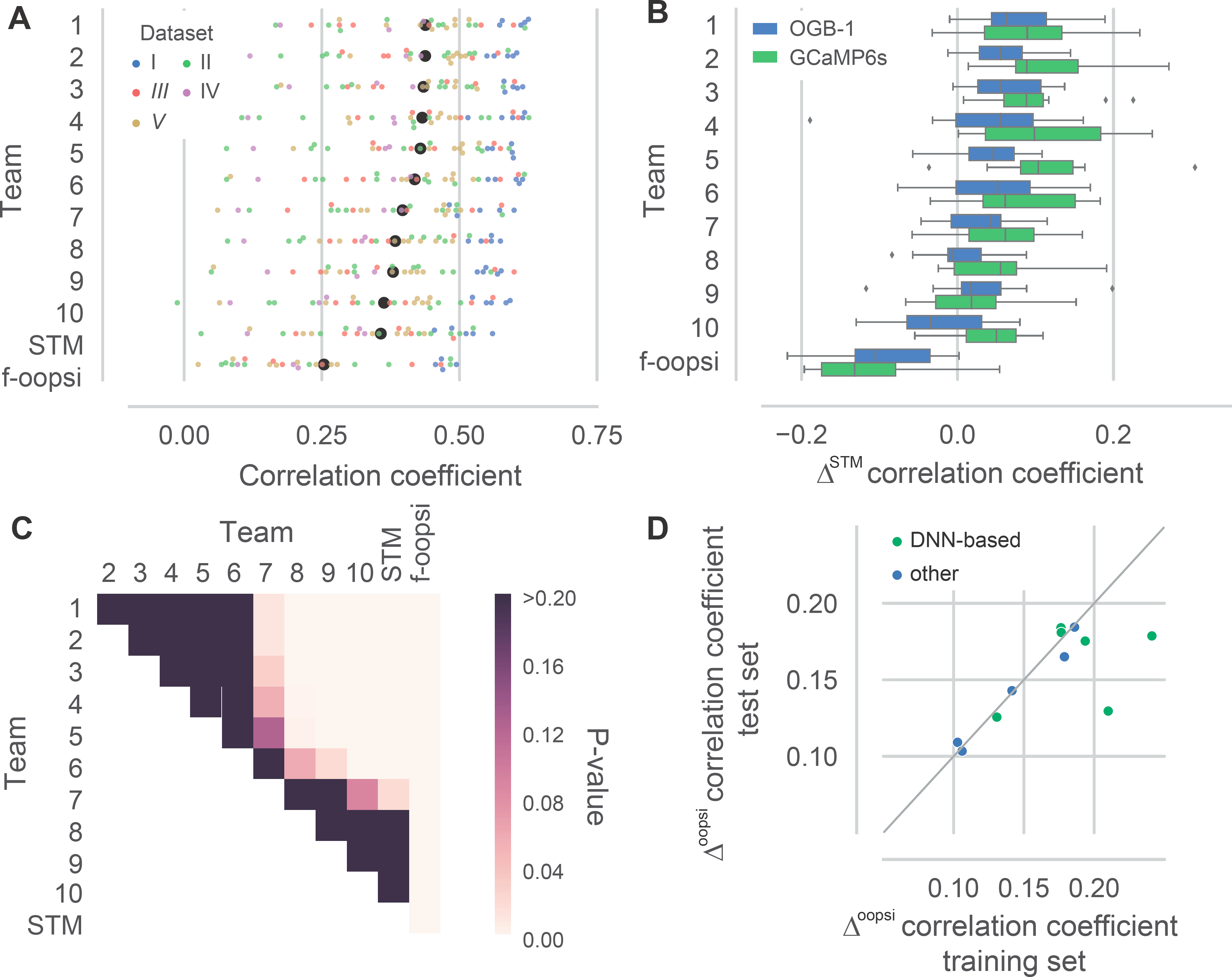
Contributed algorithms outperform state-of-the-art. A. Correlation coefficient of the spike rate predicted by the submitted algorithms (evaluated at 25 Hz, 40 ms bins) on the test set. Colors indicate different data sets (for details, see Table 1). Data sets I, II, and IV were recorded with OGB-1 as indicator, III and V with GCaMP6s. Black dots are mean correlation coefficients across all N = 32 cells in the test set. Colored dots are jittered for better visibility. STM: Spike-triggered mixture model^6^; f-oopsi: fast non-negative deconvolution^13^ (continued next page)B. Difference in correlation coefficient on the test set to the STM, split by the calcium indicator used in the data set. C. P-values for difference in mean correlation coefficient on the test set for all pairs of algorithms (Repeated measured ANOVA, N = 32 cells, main effect of algorithm: P < 0.001, shown are p-values for post-hoc pairwise comparisons, corrected using Holm-Bonferroni correction) D. Difference in correlation coefficient split by algorithm type on the training and test set, respectively, to the f-oopsi-result correcting for systematic differences between the training and the test set.

**Table 4.** Summary ofalgorithm performance. Δ correlation is computed as the mean difference in correlation coefficient compared to the STM algorithm. Δ var. exp. in % is computed as the mean relative improvement variance explained (r^2^). Note that since variance explained is a nonlinear function of correlation, algorithms can be ranked differently according to the two measures. All means are taken over *N* = 32 recordings in the test set, except for training correlation, which is computed over *N* = 60 recordings in the training set.

| Team | train correlation | test correlation | $\Delta$ correlation | $\Delta$ var. exp. % | AUC | Info |
| --- | --- | --- | --- | --- | --- | --- |
| 1 | 0.4823 | 0.4382 | 0.0810 | 44.1 | 0.846 | 2.922 |
| 2 | 0.4727 | 0.4378 | 0.0806 | 42.0 | 0.846 | 3.118 |
| 3 | 0.4730 | 0.4347 | 0.0775 | 41.8 | 0.851 | 3.085 |
| 4 | 0.5374 | 0.4325 | 0.0753 | 42.9 | 0.815 | 2.816 |
| 5 | 0.4900 | 0.4291 | 0.0719 | 40.5 | 0.842 | 2.725 |
| 6 | 0.4752 | 0.4188 | 0.0617 | 36.5 | 0.822 | 2.778 |
| 7 | 0.4379 | 0.3967 | 0.0395 | 22.2 | 0.829 | 2.797 |
| 8 | 0.5063 | 0.3833 | 0.0261 | 13.1 | 0.816 | 2.415 |
| 9 | 0.4271 | 0.3794 | 0.0222 | 13.5 | 0.815 | 2.816 |
| 10 | 0.3992 | 0.3629 | 0.0058 | 11.0 | 0.784 | 2.253 |
| STM | 0.4024 | 0.3572 |  |  | 0.821 | 2.468 |
| f-oopsi | 0.2964 | 0.2538 | -0.1010 | -40.4 | 0.658 | 1.107 |

To explore the generality of our findings, we additionally analyzed the performance of the algorithms at different temporal resolutions and using different evaluation measures. To this end, we computed the average correlation coefficient between the inferred and the true spike rates for bins of 40, 83, 167 and 333 ms, respectively (Fig. 2). As expected, the average correlation increased with increasing bin width (e.g. for algorithm by team 1: 0.44 to 0.73). Interestingly, the rank of the algorithms was consistent across bin widths. In addition, we evaluated the performance of the algorithm using the AUC and information gain (Fig. 3, Table 4, see Methods). The AUC measures the accuracy with which the presence of spiking in a given bin is detected, neglecting differences in the number of spikes. The information gain provides a model-based estimate of the amount of information about the spike rate extracted from the calcium trace ^6^. The ranking of the algorithms was broadly consistent with the ranking based on correlation, despite minor differences.

**Figure 2:**
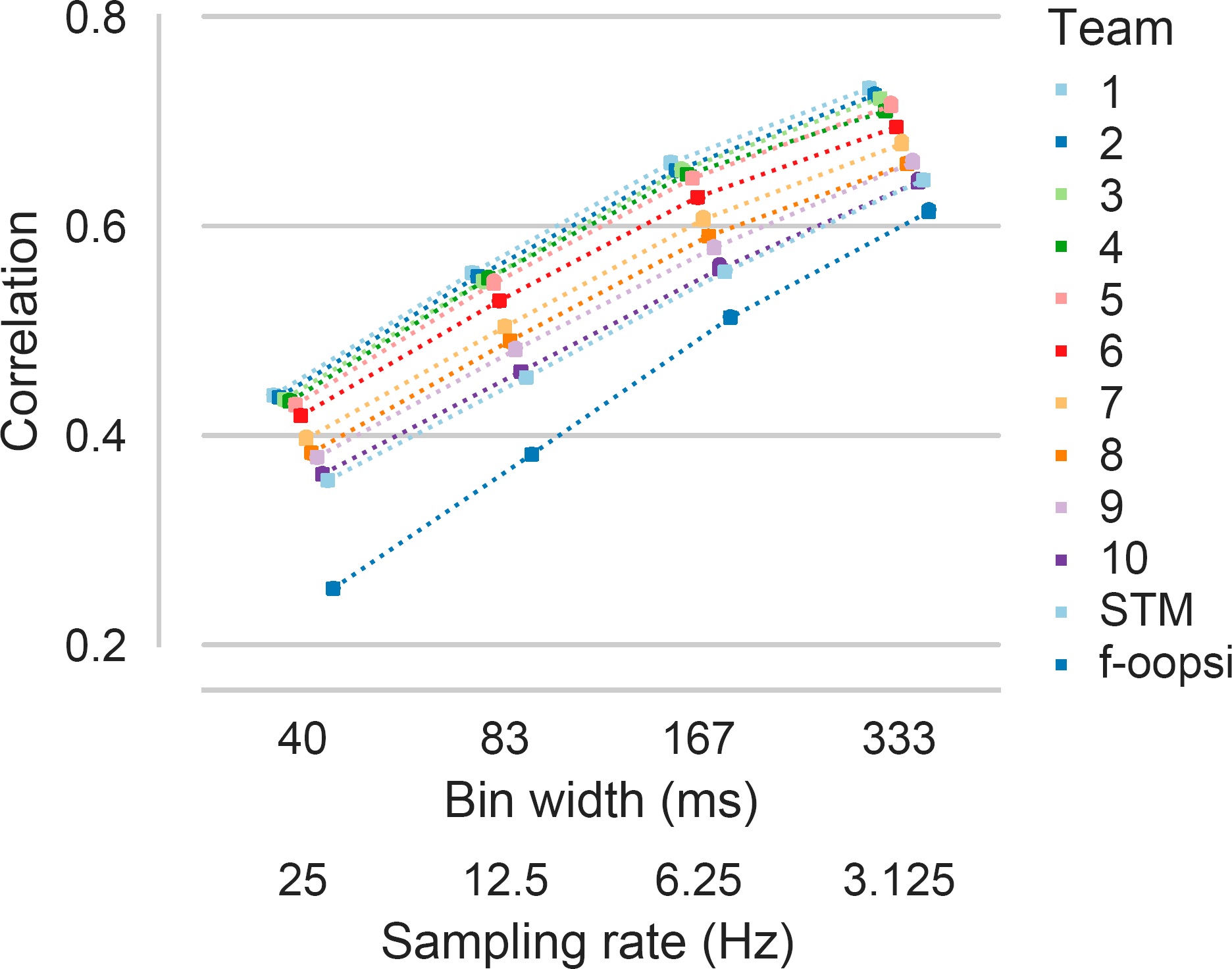
Temporal resolution does not change the ranking of algorithms. Mean correlation between inferred and true spike rates evaluated at different temporal resolution/sampling rate on all N = 32 cells in the test set. Colors indicate different algorithms. Colored dots are offset and connected for better visibility. STM: Spike-triggered mixture model^6^; f-oopsi: fast non-negative deconvolution^13^

**Figure 3:**
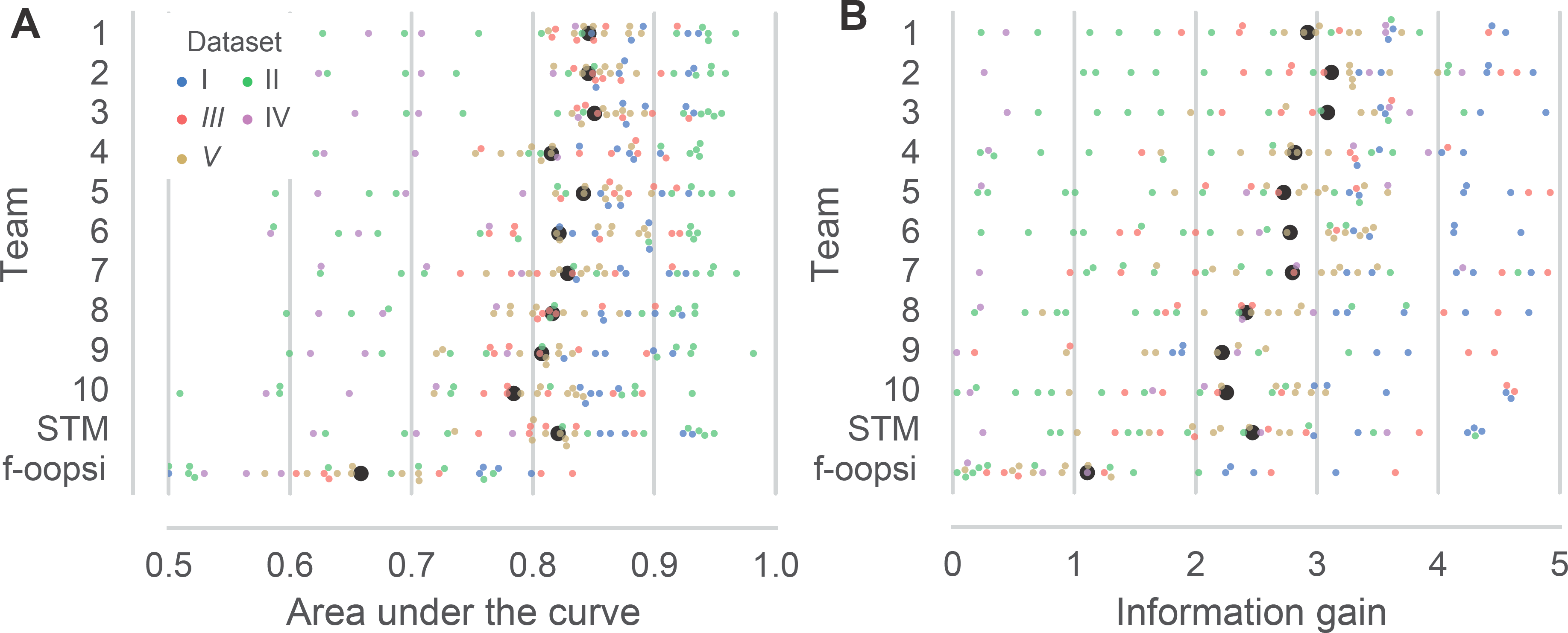
Different spike inference metrics reach similar conclusions. A. Area under the curve (AUC) of the inferred spike rate used as a binary predictor for the presence of spikes (evaluated at 25 Hz, 50 ms bins) on the test set. Colors indicate different datasets. Black dots are mean correlation coefficients across all N = 32 cells in the test set. Colored dots are jittered for better visibility. STM: Spike-triggered mixture model^6^; f-oopsi: fast non-negative deconvolution^13^ B. Information gain of the inferred spike rate about the true spike rate on the test set (evaluated at 25 Hz, 40 ms bins).

As the algorithms in the top group used a range of algorithmic strategies, we wondered whether they also made different predictions, e.g., each capturing certain aspects of the spike-calcium relationship but not others. However, the predictions of the different algorithms are typically very similar with an average pairwise correlation coefficient among the first six algorithm of 0.82 ± .04 (mean ± SD, Figure 4). Also, averaging the top six predictions in an ensembling approach did not yield substantially better performance (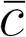 = 0.4436 compared to 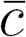 = 0.4382 for Team 1). This indicates that despite their different algorithmic strategies, all algorithms capture similar aspects of the spike-fluorescence relationship.

**Figure 4:**
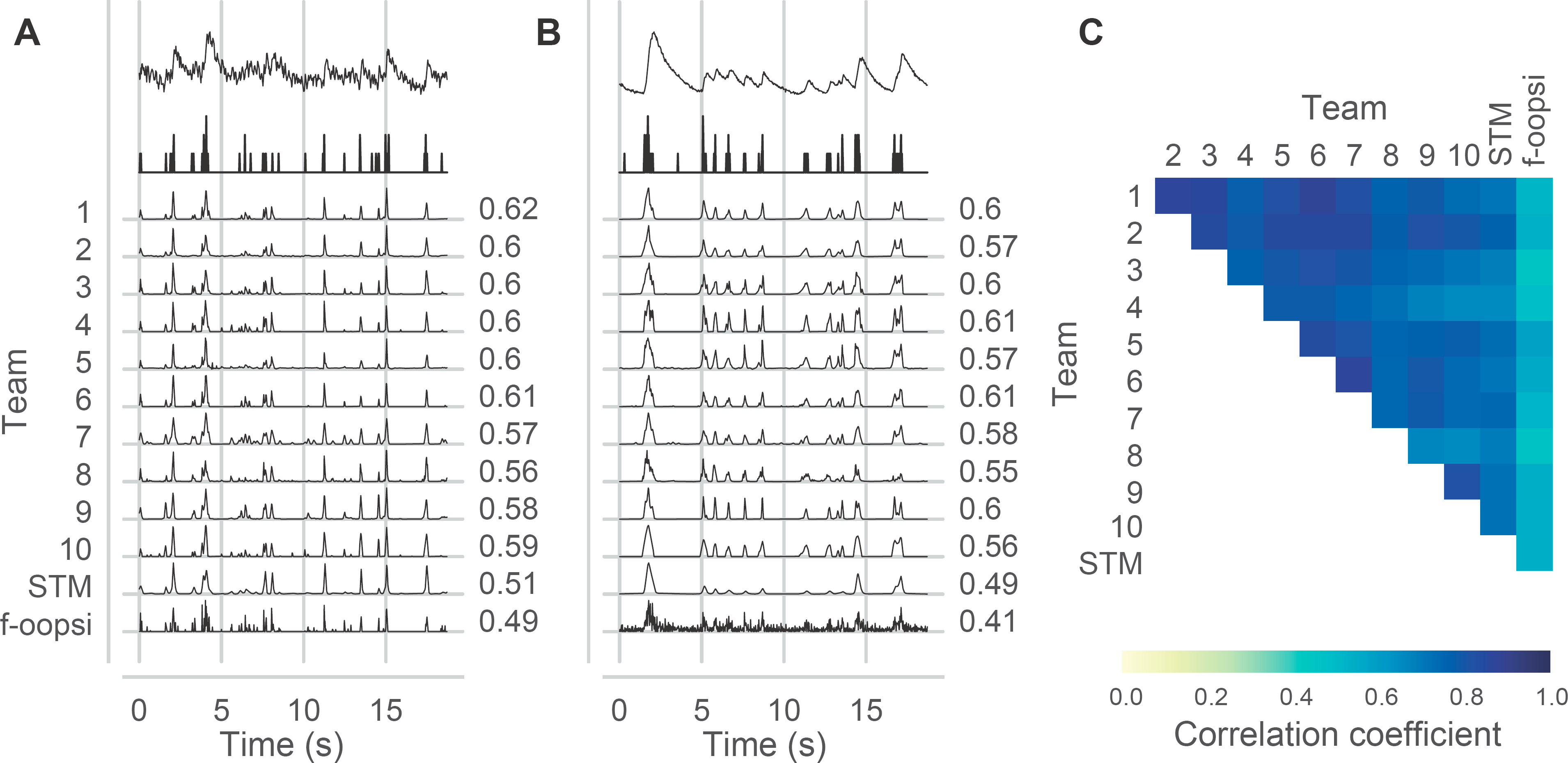
Top algorithms make highly correlated predictions. A.-B. Example cells from the test set for dataset 1 (OGB-1) and dataset 3 (GCaMP6s) show highly similar predictions between most algorithms. C. Average correlation coefficients between predictions of different algorithms across all cells in the test set at 25 Hz (40 ms bins).

## Discussion

In summary, the *spikefinder* challenge has shown that a community competition making use of suitable benchmark data can catalyze algorithmic developments in neuroscience. The challenge triggered a range of new and creative approaches towards solving the problem of spike rate inference from calcium data and improved the state-of-the-art substantially. The challenge did not distill the optimal strategy out of the different possible algorithmic approaches, something we had initially hoped for; rather, it showed that – given the current data – a range of approaches yield very similar outcomes.

### Different algorithmic strategies for spike rate inference

Interestingly, algorithms based on very different approaches yielded very similar performance. For example, algorithms based on generative models such as those by Team 1 and 6 perform on par with – in principle – more flexible deep learning-based approaches. Each algorithm comes with their own advantages and disadvantages regarding speed, interpretability, and incorporation of prior knowledge. For example, training the DNN-based models can be computationally quite costly and their efficient use may require specialized hardware such as GPUs. In practice, when a trained algorithm is applied to infer spike rates, we found all DNN-based method comparably efficient with a run time of less than a second per recording. With supervised methods, care has to be taken when using complex models to avoid overfitting the training set, as this could lead to false confidence about the prediction performance on new data. In fact, we observed quite heavy overfitting for two of the DNN-based approaches (Fig. 1D). Nevertheless, supervised spike inference algorithms have been shown to generalize well to new data sets for which no data had been used during training^6^, indicating that adapting supervised algorithms to new settings like indicators with different dynamics should be reasonably straightforward. In contrast, the algorithms based on generative models may be less easily adapted to novel settings as indicator dynamics, saturation or adaption effects and noise properties need to first be accurately assessed. In addition, inference in such models can be more time consuming as shown by the performance of the MLspike algorithm with an average of 15 seconds per recording. Hybrid approaches such as pursued here by Team 9 or more recently by ref ^23^ may offer a way towards combining the respective strengths of both approaches.

### Is spike rate inferences saturated?

The *spikefinder* challenge raises the question of what the actual performance bound of an ideal decoder is. Model simulations can help to answer these questions^7,12^, but their interpretation is limited by the accuracy of the model regarding indicator dynamics, noise structure, and other experimental factors^6^. For example, in vitro recordings zooming in on individual neurons will have a different maximal performance than recordings in awake, behaving animals. Of course, the achievable upper bound on performance always depends on the desired temporal resolution (Fig. 2) and experimental factors. For example, cells in data set I recorded at very high sampling rates using 3D AOD scanning yielded on average much higher correlation than neurons recorded using the same indicator in the same area with much lower scan rate (Fig. 1A). It remains to be seen whether new and larger data sets of simultaneously recorded and imaged neurons will yield further improvements and distinguish more clearly between different algorithmic strategies. It will also be interesting to see whether new indicators will allow for more precise spike rate inference.

### Evaluation of spike rate inference

We also considered the AUC and information gain as alternatives to our primary evaluation measure, the correlation coefficient. While the latter is easy to interpret and more sensitive than the AUC, it is still invariant under global scaling of the predicted spike rate^6^. Although information gain as a model based measured is considered a canonical model comparison criterion for probabilistic predictions^6,24^, it is considered less intuitive by some.

In general, all three measures yielded similar estimates of the ranking of the algorithms, with the AUC resolving the present differences least. In fact, different metrics can in principle lead to different conclusions about which algorithm is optimal since the metric contains part of the task specification^25^. Metrics for spike rate inference are a matter of current debate in the literature — see for example refs.^11,26^ for recent proposals.

### Design considerations for future challenges

In addition to improving on the state-of-the-art, competitions such as the *spikefinder* challenge can boost standardization of algorithms, something that has been lacking from neuroscience analysis tool chains^27^. For example, some of the preprocessing choices made for this challenge triggered a debate about the best way to handle several of the processing steps. For example, we resampled all data to 100 Hz for ease of comparison, which induced problems for some of the submitted algorithms through the properties of the used filter. In addition, most participating teams found it necessary to introduce means of adapting the model parameters to the specific data set. These differences may have been introduced through different preprocessing procedures in the labs that contributed data and even between different scanning methods and speeds within the same lab (3D AOD vs. galvo scanning vs. resonsant scanning).

Even greater care should be taken to avoid such confounds in future competitions on this topic. In particular, a future challenge should explicitly address the potential of each algorithm to easily adapt to a data set not previously seen as part of the training set, testing for the transfer learning capabilities of each algorithm. It would also be interesting to explicitly evaluate algorithms for different recording conditions (e.g. in-vitro vs. awake), as the difference in recording conditions could even make different algorithmic strategies optimal.

Finally, the challenge was performed on traces extracted from the raw imaging data by averaging all the pixels within manually placed regions-of-interest (ROIs). It is thus possible that the extracted signals contain contamination from the neuropil or were suboptimally placed, a problem tackled by methods that combine ROI placement and calcium-trace extraction in a single algorithm^5,28^. However, at least for data with simultaneous imaging and electrophysiological recordings, it is not clear how methods integrating ROI placement and spike rate extraction should be evaluated and compared to regular data, since the recording electrode is always present in the picture, adding a confound to automated ROI extraction through the different image statistics.

### Conclusion

We believe that quantitative benchmarks are an essential ingredient for progress in the field, providing a reference point for future developments and a common standard with regards to how new algorithms should be evaluated. We strongly believe that many fields of computational neuroscience can benefit from community-based challenges to assess where the field stands and how it should move forward. As for the problem of spike rate inference from two-photon imaging, the *spikefinder* challenge should not be considered the last word in this matter: More comprehensive data sets and new functional indicators may require organizing another round of community-based development, further pushing the boundaries of what is attainable. Which algorithm to choose? The answer to that depends on a lot of factors, including performance, desired programming language, envisioned run time and not the least the simplicity of the method — certainly, an algorithm consisting of ten simple lines of code like that by team 10 is more intuitive than a highly nonlinear DNN. The algorithms submitted as part of this challenge offer a range of options regarding these criteria and will provide a solid basis to further advance the field.

## Methods

### Data

The challenge was based on data sets collected for a previous benchmarking effort^6^ and the publicly available cai-1 data set from crcns.org^19^. Details about the recording region, scan method, indicators, scan rate and cell numbers are summarized in Table 1 and described in detail in Theis et al. (2016). All data was resampled to 100 Hz independent of the original sampling rate. Upon request during the challenge, we made the data available at the native sampling rate.

### Challenge organization

For the challenge, we split the available data into training and test sets (see Table 1). The training set contained both calcium and spike data, while for the test set, only calcium data was available during the challenge period. We made sure that multiple recordings from individual neurons contained in some data sets were either assigned to the training or the test set. The GENIE datasets were only used as training data, since they are completely publicly available and consist of recordings from individual zoomed-in cells.

The data and instructions were available on a dedicated website, based on an open-source web framework (https://github.com/codeneuro/spikefinder). There was a discussion board linked from the website to allow for questions and discussion among participants. Each team could make multiple submissions, but during the challenge period, only results on the training set were shown. The challenge ran from 30/11/2016 to 04/05/2017.

### Algorithms

The submitted algorithms are described in detail in the Appendix. For comparison, we used publicly available implementations of the STM algorithm^6^ and fast-oopsi^13^. STM parameters were optimized on the entire training set.

### Evaluation

The evaluation of the submissions was done in Python using Jupyter notebooks. All evaluation functions and notebooks are available at https://github.com/berenslab/spikefinder_analysis.

We used the correlation coefficient c between the inferred and the real traces resampled to 25 Hz (40 ms time bins) as primary quality measure. To make the observed increase in correlation more interpretable, we converted it to variance explained *r*^2^ and report the improvement in performance as the average increase in variance explained compared to the STM algorithm:

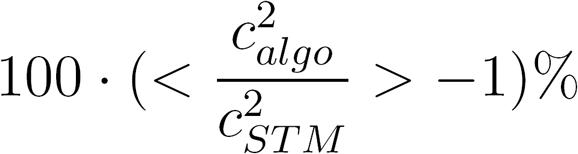

Here, <> denotes an average over cells, omitting the dependence of c on cells for clarity. For completeness, we also computed the area under the ROC curve (AUC) and the information gain as in ref. ^6^. We used the `roc_curve` function from `scikit-learn` ^29^ to compute the AUC for classifying whether or not a spike was present in a given bin. Assuming Poisson statistics, independence of spike counts in different bins, an average firing rate λ and a predicted firing rate of λ_t_ at time *t*, the expected information gain (in bits per bin) can be estimated as

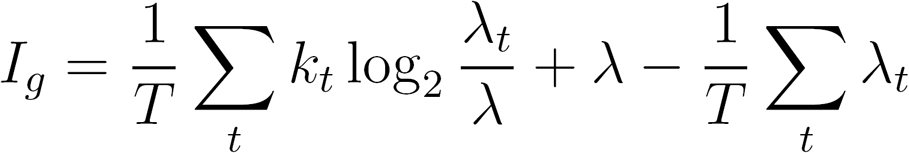

Since the different algorithms were not necessarily optimized for this model, we transformed the predicted firing rate λ_t_ using a piecewise linear monotonically increasing function *f* optimized to maximize the information gain across all cells^6^.

We used the R package `afex` to compute a repeated measures ANOVA on the correlation coefficients with within-subject factor algorithm and cells as subjects. Pairwise comparisons between algorithms were performed using the `lsmeans` package with Holm-Bonferroni correction for 66 tests.

## Acknowledgements

Awards for the *spikefinder* competition were kindly provided by Carl Zeiss AG, Oberkochen, Germany and the Bernstein Center for Computational Neuroscience, Tübingen, Germany.

## Author contributions

PB, AST and MB designed the challenge; JeF, JS, NJS provided web framework; PB and JeF ran the challenge; TD, NC, TM, AS, JHM, ST, PM, PR, SG, RF, JoF, LP, MP, KDH, TAM, DR submitted algorithms; JR, EF, TE, MRR, AST provided data; PB analyzed the results with input from LER and MB; LT preprocessed data; PB wrote the paper with input from all authors.

## Correspondence

Correspondence and requests for materials should be addressed to P.B. (:">).

## Appendix

For all algorithms, we denote the spike train *s*, the fluorescence trace ***f*** and the underlying calcium signal ***c***, where applicable. We observe a total of *T* time bins, and the measurement in time bin t is written *s_t_, f_t_* and *c_t_*, respectively.

### Team 1 — T. Deneux

The MLspike algorithm^7^ is a model-based Bayesian inference algorithm. Similarly to the method by Vogelstein et al.^4^, the conversion of neuronal spiking activity to calcium fluorescence is modeled by a biophysical dynamical system, and a two-ways filtering scheme is applied to estimate the hidden dynamics of the intracellular calcium concentration given the noisy fluorescence recording. MLspike implements two major improvements over previous models: The first one is an extension of the biophysical model including a slowly drifting baseline, which allows disentangling a wide range of noises often observed in the real data from the spike-related signals. The second one is to represent probabilities as dense arrays rather than using Monte-Carlo approximations, namely making MLspike a histogram filter instead of a particle filter, which improves both speed (at least for a models hidden state dimension not greater than 2) and accuracy.

For the spikefinder competition, MLspike was set to estimate a-posteriori probabilities 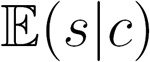 rather than maximum-a-posteriori spike trains arg max_s_ *p*(*s|c*). The biophysical model entailed a drifting baseline and nonlinear calcium to fluorescence conversion (i.e. saturation for OGB dataset; polynomial supralinearity for GCaMP6 dataset), and therefore had 6 or 7 parameters. One of these parameters (the a-priori spiking rate) was fixed while the 5 or 6 remaining ones were estimated independently for each training dataset so as to maximize the match between the estimated and observed spikes. This was preferred to using MLspikes autocalibration method on each individual neuron, because the data appeared too noisy for this autocalibration to perform accurately. The match between the true and inferred spike rate was defined as the correlation after resampling to 25 Hz between the true spike train, and the best post-processed version of the estimated spike train, were postprocessing consisted in applying an additional temporal smoothing and a time shift to account for apparent differences between data sets. Once these optimizations were performed, the same model and postprocessing parameters were applied to the test data sets. Interestingly, the hyperparameter optimization strategy pursued here was very similar to that chosen by Team 6.

Code is available at https://github.com/MLspike.

### Team 2 — N. Chenkov, T

McColgan This algorithm is based on a convolutional neural network, which receives the calcium signal and an index vector as input, denoting the data set the inputs come from. The network consists of eight convolutional layers and one recurrent layer (LSTM) (see Fig. 5A). We optimized the parameters by maximizing the Pearson correlation coefficient with the ground-truth spiking data at 25 Hz using the ‘Adam’ optimizer with 50 epochs.

The first layer consists of 10 units. Each unit uses a kernel with a width of 3 seconds (300 time steps) that is correlated with the input calcium signal. The learned kernels catch a basic repertoire of spike-related calcium dynamics (see Fig. 5A). The output of this layer is passed through a hyperbolic tangent activation function (‘tanh’). This layer is followed by multiple convolutional layers with smaller kernel widths (100 and 50 ms, or 10 and 5 time steps, respectively) and with rectified linear activation functions (‘ReLU’). The data set indicator is concatenated with the output of the second layer and feed into the input of the third layer. We observed that the data set indicator improved the performance of the model, possibly setting different states of the recurrent layer dynamics. Moreover, a bidirectional LSTM layer is fed by the input of the third layer, and its output is added to the input to the fourth layer. The following four layers have decreasing size, with the last layer consisting of a single unit.

To distribute the amount of information that different units are carrying, dropout is applied at the output of the first five convolutional layers.

Code is available at https://github.com/kleskjr/spikefinder-solution.

### Team 3 — A. Speiser, S. Turaga, J. H. Macke

We trained neural networks consisting both of convolutional and recurrent layers to learn a mapping from fluorescence trace to neural spiking. In contrast to other methods, we trained the network to approximate a correlated posterior conditional probability distribution *q(s|f)* of a spike-train *s_t_=_0_…_T_* given a fluorescence trace *ft=o…_T_*. We use a recurrent layer to model an autoregressive conditional probability distribution to account for correlations in this posterior between spike probabilities and previously sampled spikes, similar to ref ^30^. The temporal ordering over spikes is used to factorize the joint distribution as *q_ϕ_(s|f)* = ∏*_t_ q_ϕ_(s_t_|f, s_0_,…, s_t-1_)*, by conditioning spike probabilities at t on all previously sampled spikes. The resulting stochastic RNN models a correlated posterior conditional distribution over spike trains (See fig. 6). The stochastic RNN samples correlated spike trains (similar to MLSpike) which might be useful for certain applications. However, none of the performance measures used in this challenge are sensitive to this property, as they are based on marginal firing rate predictions.

**Figure 5:**
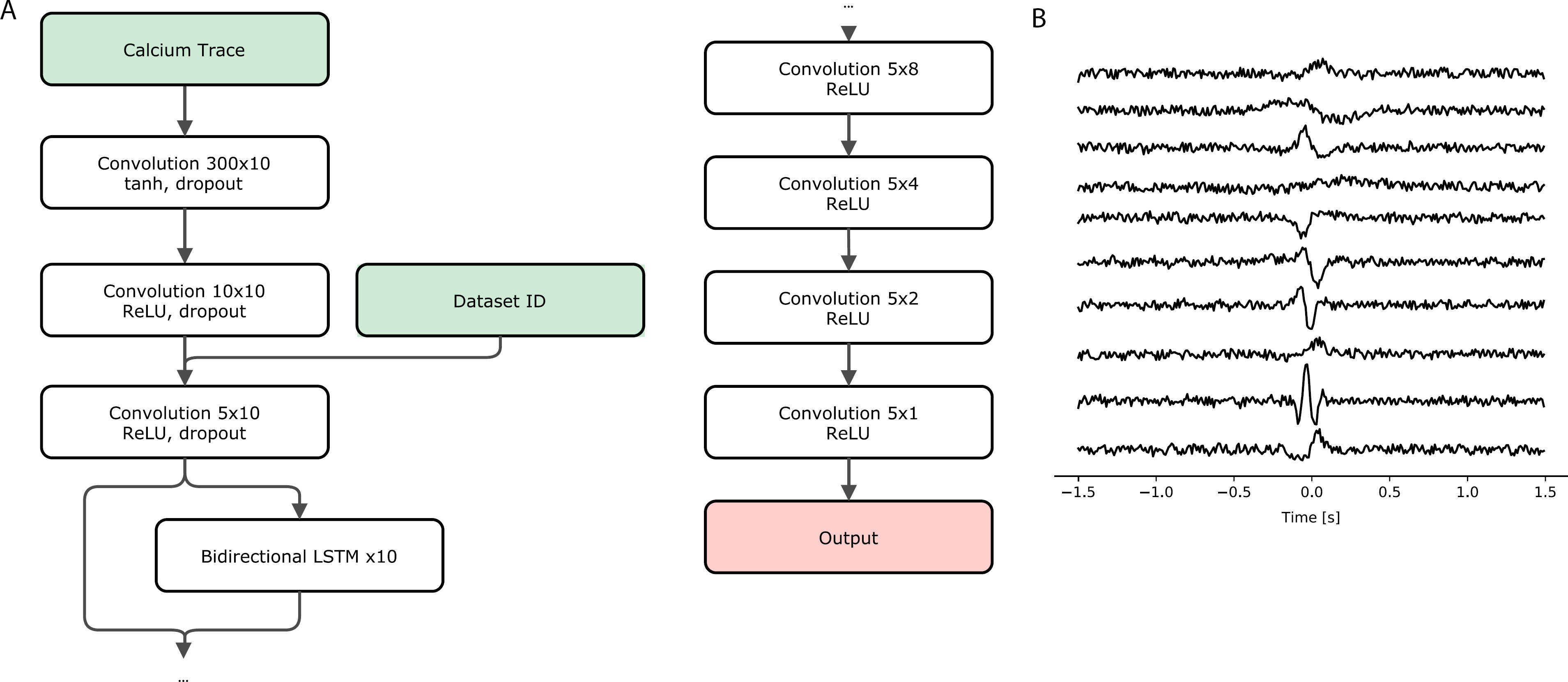
A. Network Architecture. In the convolutional layers the notation *mxn* denotes n units with kernel width of m time steps. B. Example of convolutional kernels learned by the model. The ten kernels of the first convolutional layer can describe a wide range of transient calcium dynamics.

**Figure 6:**
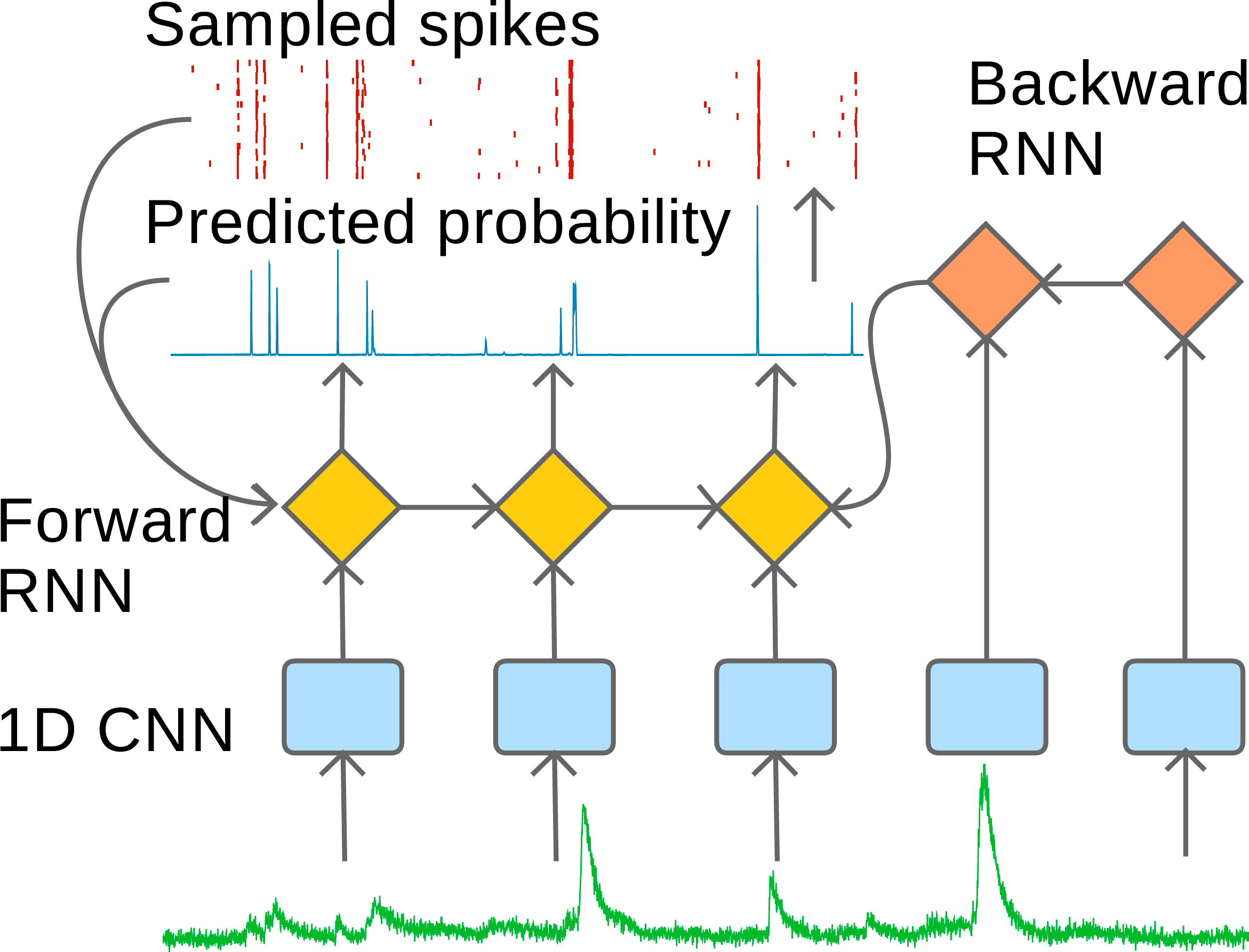
Network Architecture We use a multi-layer network architecture: Fluorescence-data is first filtered by a deep 1D convolutional network (CNN), providing input to a stochastic forward running recurrent neural network (RNN) which predicts spike-probabilities and takes previously sampled spikes as additional input. An additional deterministic RNN runs backward in time and provides further context.

As our objective function we used the binary cross entropy between our predictions and the true spike train to train the model in a supervised fashion

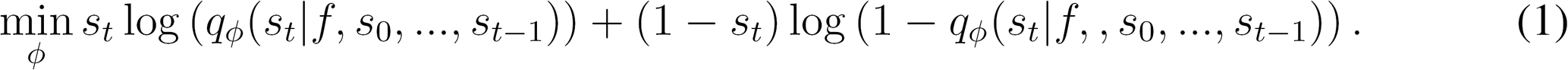

In separate work^23^, we developed an approach for training this network in an unsupervised fashion using variational auto-encoders^31^ that can perform inference on a wide range of biophysical generative models (e.g. using the ones used by team 1). For the challenge, all available training data was labeled (i.e. ground truth spikes were provided), and we therefore trained the network using supervised learning.

Our architecture contains one forward running RNN that uses a multi-layer CNN with leaky ReLUs units to extract features from the input trace. The outputs of the forward RNN and CNN are transformed into Bernoulli spike probabilities through a dense sigmoid layer. Additional input is provided by a second RNN that runs backwards and also receives input from the CNN. Forward and backward RNN have a single layer with 128 gated recurrent units each^32^. In order to generate a single marginal probability distribution *qϕ(s_t_|f)* for evaluation, samples drawn by running the stochastic RNN 50 times were averaged. We trained one separate network for each dataset.

To minimize the artifacts introduced by upsampling the data to a common imaging rate, we performed our own pre-processing (including percentile-detrending, normalizing and resampling) where we kept the fluorescence traces closer to the original recording frequency (i.e. 50, 50, 75, 12.5, 75 Hz for data sets 1-5 respectively). For the rare cases where the true spike train contains bins with multiple spikes at this rate, we clip the values to be binary. For training we split the traces into short snippets and arranged them into batches of size 10. For the rare cases where the true spike train contains bins with multiple spikes at this rate, we clip the values to be binary. Furthermore we used stochastic gradient descent with the Adam optimzer (using default parameters).

To find good hyperparameters we performed a small grid search on the following parameters and chose the best model using cross validation: learning rate {4e^-4^,1e^-3^}, number of convolutional filters per layer {20/15/15/10, 35/30/20/10}, length of trace snippets {100, 200, 300}. Performance proved to be rather robust to the exact choice of hyperparameters.

The RNN produced suboptimal results on the fourth dataset (OGB, 7.8 Hz), and we therefore used a simple factorizing CNN on this data-set, at an upsampled rate of 4x the imaging rate. The RNN architecture achieved a correlation coefficient of 0.417 on the validation set against 0.455 when using the CNN.

Code is available at https://github.com/mackelab/DeepSpike.

### Team 4 — P. Mineault

This submission casts the problem as a supervised learning problem, where the goal is to estimate the parameters of a basis transformation *g* and an output non-linearity h such that a loss *L(μ_t_,s_t_)* between spike train *s_t_* and prediction *μ_t_* is minimized. *g* is given by a deep convolutional artificial neural network. The first layer of this network is a standard convolutional layer followed by a rectified linear (ReLU) nonlinearity^33^, mapping each calcium time series *f_t_* to 32 parallel time series indexed by *k*.

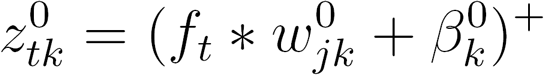

Here (x)^+^ = max(0, *x*) is the ReLU nonlinearity. We use a large window (33 time points, or 330 ms), batch normalization, as well as a dropout fraction of .3. This initial layer is followed by seven adjustment layers in the style of a residual network ^34:^

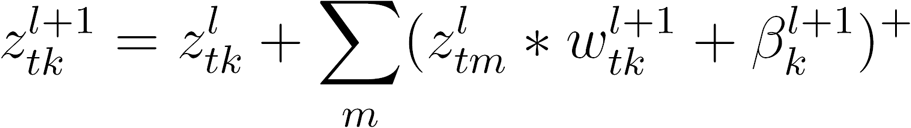

These adjustment layers use smaller windows (9 time points, or 90 ms). The nonlinear component of each layer is batch normalized. Finally, the output is composed linearly via:

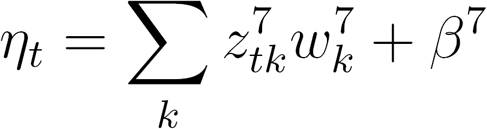

The output nonlinearity *h* was given by a ReLU nonlinearity, such that 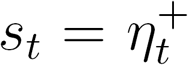. We minimized a scaled sum of squared error criterion:

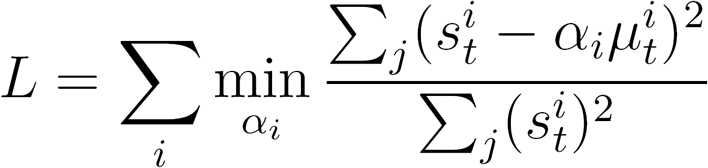

Here, *i* indexes different neurons and α*_i_* is a set of scalars which are learned alongside the other parameters of the model (*w* and *β*). One can show that the loss is equivalent to 1 — *ρ*^2^, where *ρ*^2^ is the square of the cosine similarity between prediction and spike train. The model was specified and fit using the tf.contrib.learn library in TensorFlow^35^. Model parameters were initialized with the Xavier method and fit using the Adam optimizer. One large model for all 10 recordings from the training set was fit (173 neurons) in this phase and goodness-of-fit was monitored on a leave-aside validation set to control overfitting by early stopping. Convergence took close to 200,000 iterations.

The model described so far uses fixed filters for each recording, and uses local information (≈ 1 second of data) to estimate spikes from calcium traces. To adapt filters, we learned long-range features with an unsupervised mixture density network^36 37^. The model, a 3-layer recursive neural network with 512 long-short-term memory (LSTM) nodes, was fit to the calcium data to obtain one 1536-dimensional latent vector per mini-batch, which was reduced to 32 dimensions by PCA after z-scoring. These features were processed by two fully connected layers to produce 4 hidden features *γ*_tp_. These 4 hidden features were used to additively adapt the filters 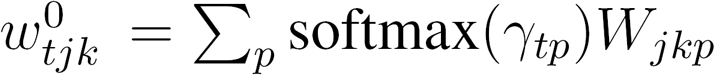 in a manner similar to attention models.

Originally, one large model was fit for all recordings. We then created refined versions of this model for each of the 10 data sets by a transfer learning process. We took the large model with its learned parameters and ran up to 50,000 extra iterations of gradient descent on just the data from the *k*th dataset.

Code is available at

https://github.com/patrickmineault/spikefinder_submission

### Team 5 — P. Rupprecht, S. Gerhard, R. W. Friedrich

In order to infer a spike probability for time bin *t* (Fig. 7A), the calcium trace located around t was used, including 25% before and 75% after *t*, totaling to 128 samples, *i.e.,* 1.28 sec (Fig. 7C). A convolutional neural network was trained to use these 128-wide windows to predict the corresponding spiking probability. To facilitate gradient ascent, we smoothed the discrete spiking ground truth with a Gaussian filter *(σ* = *√2* samples, Fig. 7B).

**Figure 7:**
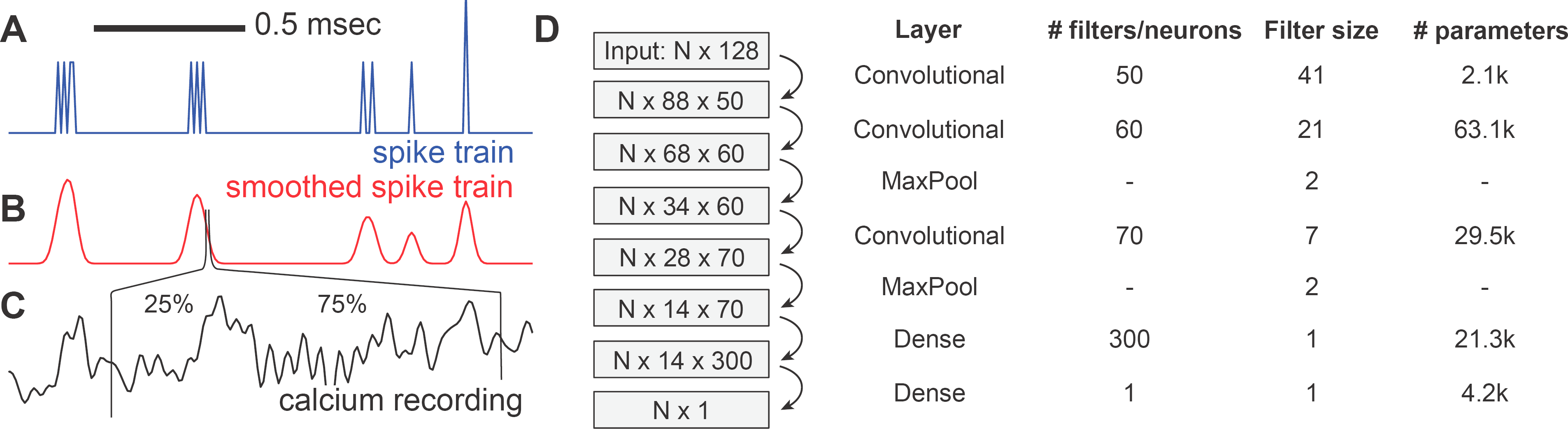
The basic convolutional neural network. A,B. The spiking ground truth was smoothed to facilitate gradient descent. C. A 1.28 sec time window of the calcium trace was used to infer the spiking probability for each time point. D. The N inputs were transformed using a neural network with three convolutional layers. The numbers in the boxes indicate the output size of the respective layers.

We implemented the convolutional neural network in Python using Keras^38^ with the Tensorflow^35^ backend (see Fig. 7 for the network architecture). The convolutional filter size, particularly for the first layer, was chosen rather large, since simple CNNs with 3 or 4 convolutional layers with small input filter sizes (3, 5 or 7) performed poorly. No zero-padding was used. The numbers of filters were chosen to increase with depth in order to allow for a larger capacity to represent higher-order features. Standard *ReLU* activation units were used after each convolutional and dense layer, except for the last dense layer, where a linear activation was used to allow the output of continuous spiking probabilities.

All parameters were chosen based on intuition gained through a small exploratory hyperparameter study using diverse 3- and 4-layer CNNs with varying filter sizes, filter numbers and input window sizes. Overfitting was controlled by randomly omitting single neurons from the training data and checking predictive performance of the CNN model for the respective omitted neuron.

Although the above-described CNN performed well when it came to fitting single datasets of the ground truth, one single model trained on all datasets usually performed not as well for any of the datasets as the same CNN trained on the respective dataset alone. To better understand this, it was quantified how well a model that had been fitted to predict spikes for neuron *i* can make the same kind of predictions for neuron *j*. To this end, a low-capacity CNN (with two locally connected convolutional layers and one dense layer) was fitted for each neuron i. The small size of the network together with a high dropout rate during training (50% after each layer) was used to prevent overfitting. This model was then applied to predict spiking probabilities both for neuron i and all neurons *j ≠ i*, resulting in a matrix of ‘predictive power’ (measured with the Pearson correlation coefficient between prediction and ground truth, identical to the evaluation of the spikefinder competition computed (Fig. 8A). For instance, row 55 shows how well spikes of neuron 55 can be predicted by the networks generated by all other neurons. Column 55, on the other hand, shows how well the model generated by neuron 55 can predict spikes of other neurons. The 5% neurons that were worst at predicting their own spiking were discarded from the following modeling, assuming bad recording quality that is not suited for inclusion into a training dataset.

**Figure 8:**
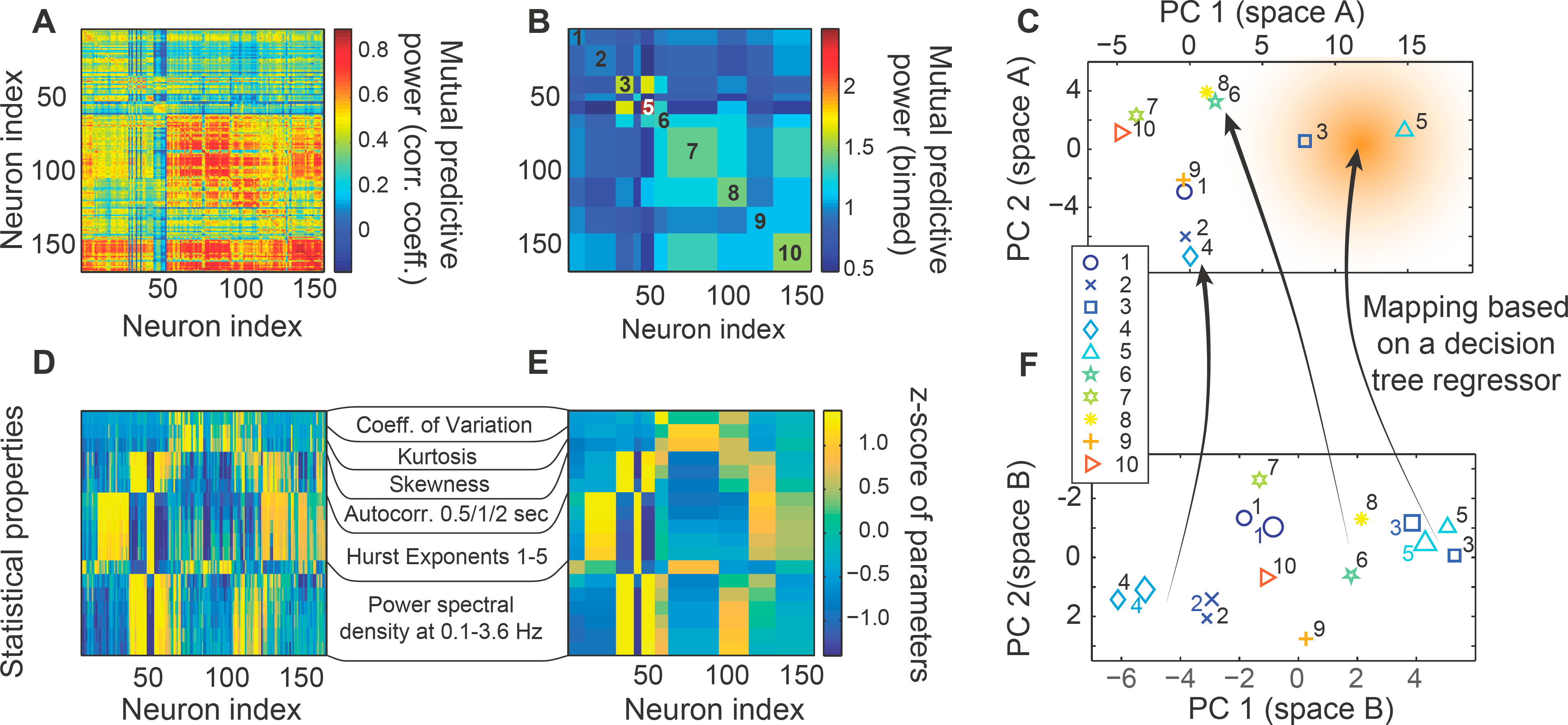
Using embedding spaces to choose datasets for focused retraining. A. Matrix of mutual predictive power, measured using the Pearson correlation coefficient between prediction and ground truth. B. Same as (A), but normalized for columns and binned to datasets. C. Principal component analysis applied to (B), keeping the first two PCs. D. Statistical parameters quantified for single neurons, standardized. E. Same as (D), but binned for datasets. F. 2D principal component space generated using (E). Symbols with numbers to the right are from training datasets, used to span the PCA space; symbols with numbers to the left are from the test dataset and were projected into the PCA space.

Normalization over columns, symmetrization of the matrix and averaging over datasets yields a matrix of predictive power, i.e., a matrix of proximity in prediction-space between datasets (Fig. 8B). A PCA of this matrix results in an embedding space that was limited to two dimensions due to the low number of datasets. Datasets close to each other in the embedding space (e.g. 2 and 4) can predict each other’s spikes very well, whereas datasets distant from each other in space (e.g. datasets 4 and 5) fail to do so. The idea behind this approach is very similar to the embedding spaces used by Team 8.

Using this approach, it is however not yet possible to map a neuron of a new dataset of unknown properties onto the right location of the embedding space above. To solve this problem, the following statistical properties of the raw calcium time traces were calculated (fig. 8D), in an approach that is similar to the long-range features of calcium traces used by Team 4:

- coefficient of variation, kurtosis, skewness
- autocorrelation of the calcium time trace with its future value in 0.5, 1 and 2 seconds
- generalized Hurst exponents of order 1-5
- the power spectral density at different frequencies between 0.1 and 3.6 Hz

We did not attempt to find a minimal set of predictive properties to reduce computation time here, but used dimensional reduction techniques to automatically extract the relevant independent components. After averaging the standardized values over datasets (Fig. 8E), we used the two first principal components to generate a map of proximity in statistical property space (Fig. 8F). This map was generated using the training datasets (numbers located on the right side of the symbols). Test datasets were mapped into this PCA space (numbers on the left side of the symbols).

To generate a mapping between the locations of the datasets in the two embedding spaces,a simple regressor *(DecisionTreeRegressor* from the scikit-learn package^29^) was fit to the training datasets (schematic arrows in Fig. 8C,F). We then used this mapping to determine the position of the test datasets in the embedding space of mutual predictive power.

Once the position in the embedding space is known for a dataset, the model that had been trained before on all datasets is retrained, but preferentially with neurons from datasets that lie close to the position in the embedding space. This preference was weighted with a function that decays exponentially over distance in the embedding space, as indicated by the red shading (Fig. 8C). Again, the functional form of the decay and the decay constant have been chosen heuristically without systematic optimization, since our goal was to showcase the power of our embedding space approach rather than finding a global optimum.

Embedding spaces as a visual and explicit intermediate step for model refinement are more easily accessible for users, allow the use of relatively small convolutional neuronal networks and can highlight similarities and differences between datasets. For example, it is interesting to see that in both embedding spaces, datasets 3 and 5 cluster together, whereas dataset 8, which uses the same calcium indicator (GCaMP6s) in the same brain region (V1), is in proximity of dataset 6 (GCaMP5k in V1). It was also observed that the datasets that use OGB-1 as indicator (1,2 and 4) tend to occupy similar regions of the embedding spaces.

This indicates that model selection is not only based on the calcium indicator and the brain region, but on hidden parameters, *e.g.,* signal-to-noise of the calcium recording, sampling rate, spike rate, temperature, indicator concentration, or others. To reliably comprise these possible hidden parameters with embedding spaces, it will be necessary to increase the number of datasets in order to support as many possible types of datasets as possible. However, the unknown dimensionality of this hidden parameter space makes it difficult to predict how many datasets would be required.

Code is available at

https://github.com/PTRRupprecht/Spikefinder-Elephant.

### Team 6 — J. Friedrich, L. Paninski

This algorithm approximates the calcium concentration dynamics *c* using a stable autoregressive process of order *p* (AR(*p*)).

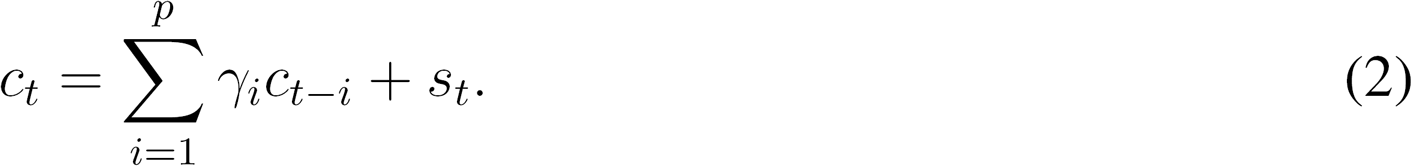

The observed fluorescence 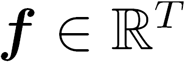 is related to the calcium concentration as^13^:

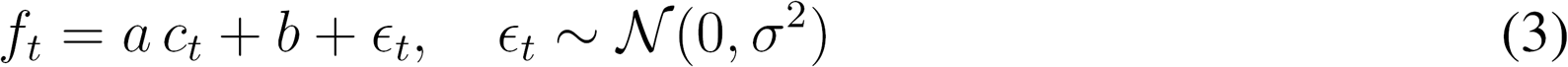

where *a* is a non-negative scalar, *b* is a scalar offset parameter, and the noise is assumed to be i.i.d. zero mean Gaussian with variance *σ^2^.* We assume units such that *a* =1 without loss of generality.

The goal of calcium deconvolution is to extract an estimate ŝ of the neural activity s from the vector of observations ***f***. This leads to the following non-negative LASSO problem for estimating the calcium concentration^8 13^:

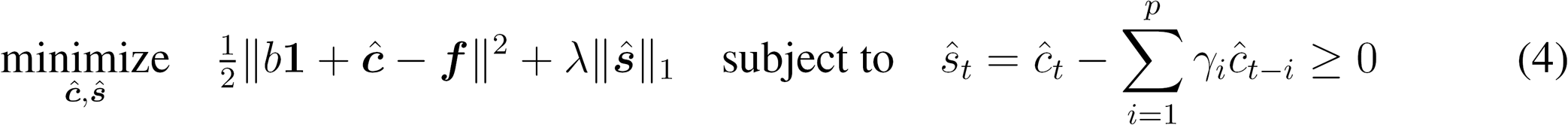

where the *l_1_* penalty on ŝ enforces sparsity of the neural activity. Note that the spike signal s is relaxed from non-negative integers to arbitrary non-negative values^13^.

Problem (4) could be solved using generic convex program solvers, however, it is much faster to use OASIS ^20^, a dual active set method that generalizes the pool adjacent violators algorithm, a classic algorithm for isotonic regression ^39^. The dual active set method yields an exact solution of Eq. (4) for p = 1 and merely a greedy one for *p* ≥ 2. Although an exact solution for the latter can be obtained by the primal active set method^8^, here p =2 is used and the greedy but faster dual method which yielded similar scores (i.e. correlation values with ground truth).

The noise level *σ* is typically well estimated from the power spectral density (PSD) of ***f*** ^5^.The parameters *γ_i_* are either known a priori for a given calcium indicator or estimated from the autocovariance function of ***f***, and possibly improved by fitting them directly. The sparsity parameter λ can be chosen implicitly by inclusion of the residual sum of squares (RSS) as a hard constraint and not as a penalty term in the objective function^5,20^. The dual problem

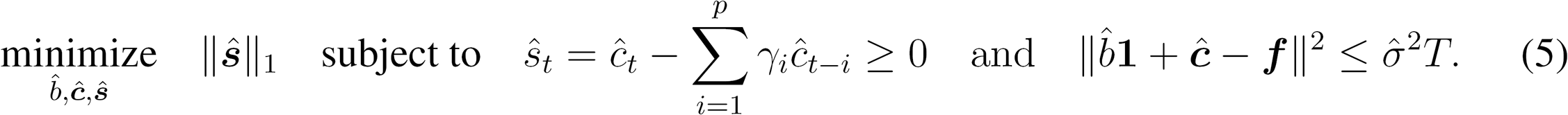

is solved by iterative warm-started runs of OASIS to solve Eq. (4) while adjusting λ, 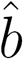 (and optionally also *γ_i_* between runs until Eq. (5) holds. We refer the reader to ^8^ for the full algorithmic details.

The above parameter choices rely on a robust noise estimate 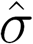 The resampling of each spikefinder dataset to a fixed frame rate introduced artifacts into the data that corrupted the autocovariance and PSD such that it was not possible to obtain reliable noise and AR estimates based on the preprocessed data. Therefore, these parameters for baseline, sparsity and AR dynamics were determined based on the training data sets and kept fix for each test trace, thus not accounting for differences between neurons within one data set. Six parameters were fit: the percentile value and window length to estimate the baseline using a running percentile, the two AR coefficients, and the slope and offset of a linear function that determines the sparsity parameter λ as function of the noise. The latter was estimated on traces that were decimated by a factor of 10 to counteract the artifacts that had been introduced by upsampling the raw data.

Running OASIS with the known parameters directly yields an estimate ŝ of the neural activity. This estimate was already good for datasets 6-10, but noticeably improved for the first 5 datasets by convolving it with some kernel *k*, to obtain the final estimate 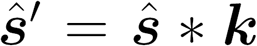. The kernel adjusts for mismatches between the actual calcium response kernel and the AR(2) model, smoothes the estimate, and accounts for the uncertainty of the exact spike timings by distributing spikes as spike rates over a few time bins. We used a kernel width of 30 bins and obtained it by averaging the closed form solutions of the least squares linear regression problem 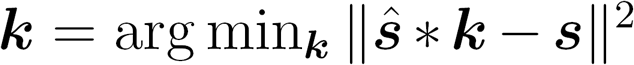 for each true spike train *s* in the training set. Interestingly, the strategy used for hyperparameter optimization used here was very similar to that used by Team 1.

Because the evaluation criterion was correlation not the residual sum of squares, we considered to further optimize the kernel for this specific criterion using gradient decent initialized at the least squares solution; however, we did not obtain significant improvements.

Code is available at

https://github.com/j-friedrich/spikefinder_submission

### Team 7 — M. Pachitariu, K. D. Harris

This algorithm has been developed as part of Suite2p, a complete calcium processing pipeline ^28^. This algorithm is called L0 deconvolution and consists of solving the following problem

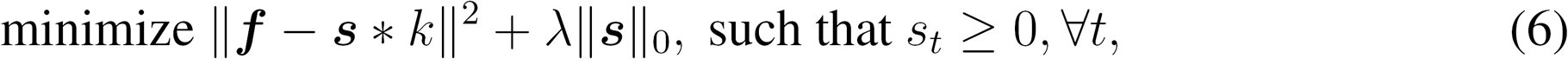

where k is the calcium kernel (assumed, or estimated), s * k is the convolution of s and *k,* and ||s||_0_ is the L_0_ norm of *s*, in other words the number of non-zero entries in s. We do not describe here the inference method for this model, but point the reader to our original derivation in ^11^. This solution is approximate, due to its greedy nature. Exact solutions have been obtained by ^40^, in the case where the positivity constraint on *s* was removed, and the calcium kernel was restricted to be exponential.

The L_0_ deconvolution model was developed as an alternative to the L_1_-deconvolution model ^5^,^8,13^. We originally believed that an L_0_ penalty would better account for the binary nature of spike trains, and allow the algorithm to return sparse spike trains. The algorithm indeed returns very sparse descriptions of the calcium data, which can deceptively look like electrophysiologically recorded spike trains. However, neither the L_0_ nor the L_1_ penalties are necessary or desirable for achieving best performance, the positivity constraint is sufficient^11^.

Here, the training data was not used to set the parameters for the deconvolution (with the exception of time lags, see below). Instead, calcium kernels were chosen to be exponentials with timescales obtained from the literature for each specific sensor^3^. Following deconvolution, dataset-specific timelags were introduced for some of the spikefinder datasets. Also, the output was smoothed with a Gaussian kernel of a preset standard deviation (80ms for spikefinder data sets, 20 ms for GENIE datasets).

Code is available at https://github.com/cortex-lab/Suite2P.

### Team 8 — B. Bolte

The algorithm used for this submission consisted of a series of stacked convolutional neural networks with filter lengths of 10 to 100 milliseconds. The model was trained to maximize the Pearson correlation between the spike probabilities predicted by the model and the ground truth spike data. To capture the non-linear dynamic characteristic of this problem, additional features were added besides the raw calcium trace, including the first and second order derivatives, as well as quadratic features. Additionally, average pooling over convolutional filters was used to capture dynamics at multiple time scales.

During experimentation, it was observed that the spike behavior varied quite a bit in different data sets. From this observation, it was inferred that different convolutional filters would perform well for different data sets. To implement this idea, “data set embeddings” were used to weight

Intuitively, these vectors represent embeddings for each data set, and the similarity between two embeddings represents the similarity of the spike behavior in each data set, since a model trained to infer spikes in the two data set would employ similar convolutional filters. This is illustrated in Figure 9.

**Figure 9:**
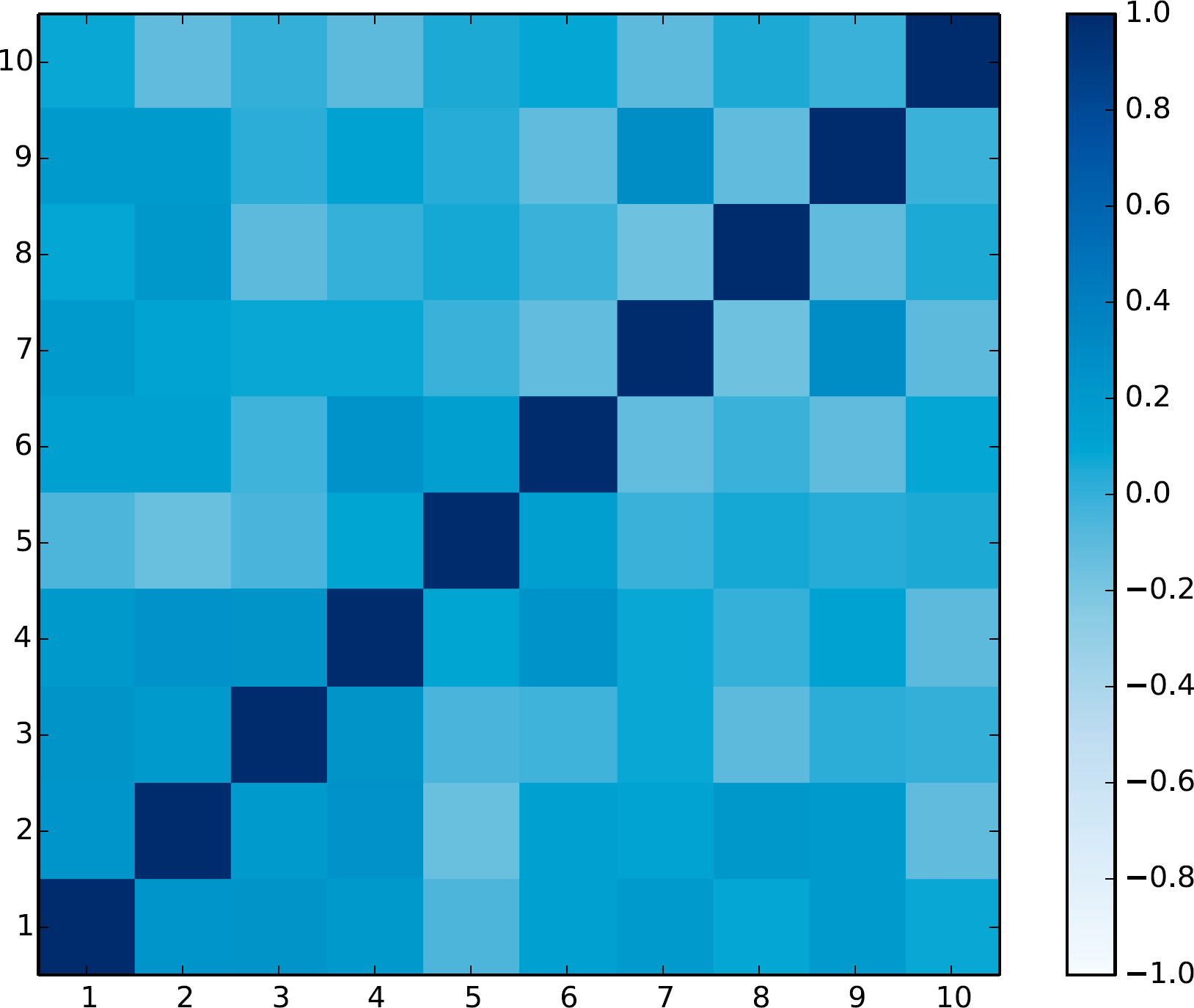
For each of the 10 data sets (5 spikefinder data sets and 5 GENIE-data sets), a unique embedding was learned, which was used to weight the output of the first convolutional layer of the network. Each cell represents the cosine similarity between the embeddings learned for that data set pair. A high similarity indicates that the two data sets used similar filters for inferring spike. the output of each convolutional filter during learning. A unique vector was learned for each data set, where the number of dimensions in the vector corresponded to the number of convolutional filters in the first layer of the model. The output of the first convolutional filter was weighted by it’s corresponding vector.

### Team 9 — T. Machado, L. Paninski

The inference framework developed by Team 9 consists of two parts: a linear encoding model that takes in spikes and outputs simulated fluorescence traces (trained on paired spike train and fluorescence data), and a simple convolutional neural network to serve as a decoding model that outputs estimates of spikes given fluorescence observations and encoding model parameter estimates. This network is trained on large data sets simulated from the encoder model. The advantage of this approach is that we can train the decoder model to “saturation” by providing it as much training data as necessary to achieve good performance. On linear-Gaussian simulated data, the neural network decoder performed comparably to OASIS^8^, a state-of-the-art inference method for efficiently efficiently solving the spike inference problem under linear-Gaussian assumptions (though both are fast enough to support online data analysis, OASIS runs significantly faster than the neural network decoder at test time).

Instead of directly using the *spikefinder* data sets to train a decoding algorithm, we generated simulated training data sets consisting of 5,000 traces, each of length 3,000 time steps. Each fluorescence trace was generated with the following second-order autoregressive model (*p* = 2):

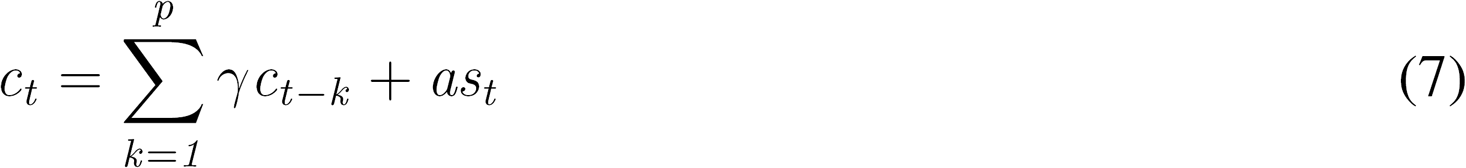

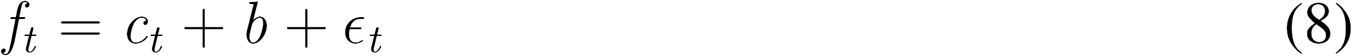

The parameterization of each trace generated using Equation 8 was random. The noise was modeled as 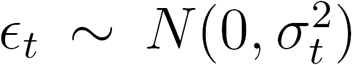 The jump size *a* of each spike was randomly sampled from a uniform distribution between 0.5 and 1.5. Each simulated spike train, *s*, was sampled from a homogeneous Poisson process with a mean firing rate between 0 and 2 Hz. The baseline drift component, *b*, was modeled by low-pass filtering white noise. The baseline drift component significantly improved inference quality, in agreement with the observation of Team 1. In contrast, the statistics of the simulated spike trains, as well as the randomized jump sizes following each spike, had a much smaller impact on decoder performance.

The time constants of the autoregressive model, *γ*, and the scale of the noise, *σ* were also sampled from random uniform distributions (such that a single decoder trained on these data could work well across multiple indicators and frame rates), but the precise parameterization was varied between model training sessions. However, in all cases, we found that a wide range of *γ* values could be learned by a single decoder model. For instance, we successfully trained decoder models across data with *γ* values spanning approximate decay times for fast OGB data recorded at 25 Hz, to slow GCaMP6S data recorded at 100 Hz. This shows that a single decoding model trained on simulated data can be used to analyze data produced by many different indicators and many different acquisition rates in agreement with Theis et al.^6^. Similarly, wide ranges of *σ* values could be learned by single decoders. However, there was a slight performance enhancement seen by training single encoders on mostly low SNR or high SNR data to improve performance on low SNR and high SNR real data, respectively.

Finally, in almost all cases, the second-order models (i.e. p =2 in Equation 7) outperformed first-order models (*p* = 1). An exception were the OGB-1 data sets, as the sensor displays very rapid rise time kinetics following action potentials and has been previously shown to be especially well-described by first-order models^13^.

To estimate the most likely spike train underlying a given fluorescence trace, we built a convolutional neural network. During training, the network was presented with f as well as parameter estimates for *σ* and *γ* given by methods published in earlier work^5^. Because the *spikefinder* data was upsampled from its native resolution and this introduced artifacts in the power spectrum of each fluorescence trace, we decimated each trace by a factor of 7-10 (depending on the approximate native time resolution of each dataset) before performing subsequent parameter fitting and analysis. The target of the network during training was the set of simulated spike trains, *s*, used to generate f using Equation 7.

A fairly simple architecture inspired by research into the construction of generative models for audio data was found to be effective ^41^. In brief, the network consists of four 1D dilated convolutional layers containing 100 units, a filter size of 32, and rectified-linear (relu) nonlinearities. The first layer was dilated by a factor of 1, the second layer by 2, the third by 4 and the fourth by a factor of 8. Dropout (rate = 0.5) was also used at each layer. Finally, a fifth 1D convolutional layer with one unit, a filter size of one, and a relu nonlinearity was used to read-out a non-negative estimate of s from each f vector provided.

This architecture contained about 950,000 parameters and could be trained on a simulated data set of 5,000 traces in about 20 minutes (over 20 epochs) using the Google ML Engine. A single model trained on simulated data that spanned a wide range of *σ* and *γ* values performed well, but an ensemble of four models, each trained on a slightly different simulated data set, worked even better-as some decoders tended to work better or worse on each *spikefinder* data set. For our submission, we chose the decoding model that worked best for each dataset to use as our submission. For data set 5, which had high firing rates, we found that convolving the results with a small Gaussian kernel resulted in a modest improvement to our inference quality.

Code is available at

https://bitbucket.org/tamachado/encoder-decoder

### Team 10 — D. Ringach

This algorithm consists of a simple linear filter followed by a static-nonlinearity *f (t) = ϕ(h(t) * s(t))* 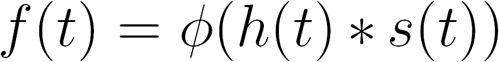. The filter h(t) is a linear combination of an even filter, estimating the mean of the signal at time t, and an odd filter, estimating the derivative of the signal at time *t*.

The even filter is a Gaussian,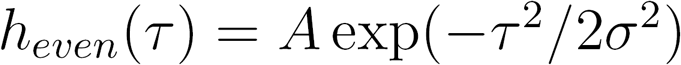, and the odd filter is the derivative of a Gaussian 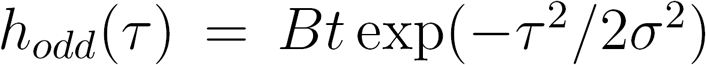. The constants A and B are such that the norm of the filters is normalized to one, ||A|| = ||B|| = 1. These two filters are linearly combined while keeping the norm of resulting filter equal to one, 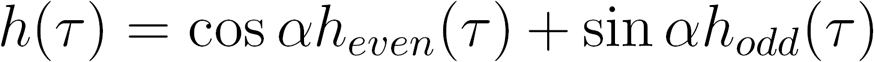. The output nonlinearity is a rectifier to a power, 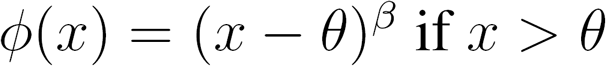, and zero otherwise.

The model has only 4 parameters, *σ, α, θ, β.* The amount of smoothing of the signal is controlled by *σ* the shape of the filter is controlled by *α,* and the threshold *θ* and power *β* determine the shape of the nonlinearity. The model is fit by finding the optimal values of *σ, α, θ, β* that maximize the correlation between its output 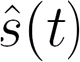 and the recorded spiking of the neuron. Matlabs `fminsearch` was used to perform this optimization, which was typically finished in about 60 sec or less for most data sets. The only pre-processing done was a z-scoring of the raw signals. In one dataset (dataset 5, GCaMP6s in V1), an extra-delay parameter between the signal and the prediction was allowed.

Code is available at https://github.com/darioringach/Vanilla.

## Footnotes

1 Notebooks and code showing how to run the individual algorithms are available at https://github.com/berenslab/spikefinder_analysis.

## References

1. Kerr, J. N. & Denk, W. Imaging in vivo: watching the brain in action. Nature Reviews Neuroscience 9, 195–205 (2008).

2. Peron, S., Chen, T.-W. & Svoboda, K. Comprehensive imaging of cortical networks. Current Opinion in Neurobiology 32, 115–123 (2015).

3. Chen, T.-W. et al. Ultrasensitive fluorescent proteins for imaging neuronal activity. Nature 499, 295–300 (2013).

4. Vogelstein, J. T. et al. Spike inference from calcium imaging using sequential Monte Carlo methods. Biophysical journal 97, 636–55 (2009).

5. Pnevmatikakis, E. A. et al. Simultaneous denoising, deconvolution, and demixing of calcium imaging data. Neuron 89, 285–299 (2016).

6. Theis, L. et al. Benchmarking Spike Rate Inference in Population Calcium Imaging. Neuron 90, 471–482 (2016).

7. Deneux, T. et al. Accurate spike estimation from noisy calcium signals for ultrafast threedimensional imaging of large neuronal populations in vivo. Nature Communications 7, 12190 (2016).

8. Friedrich, J., Zhou, P. & Paninski, L. Fast online deconvolution of calcium imaging data. PLoS Comput Biol 13, e1005423 (2017).

9. Cotton, R. J., Froudarakis, E., Storer, P., Saggau, P. & Tolias, A. S. Three-dimensional mapping of microcircuit correlation structure. Frontiers in neural circuits 7, 151 (2013).

10. Grewe, B. F., Langer, D., Kasper, H., Kampa, B. M. & Helmchen, F. High-speed in vivo calcium imaging reveals neuronal network activity with near-millisecond precision. Nature methods 7, 399 (2010).

11. Pachitariu, M., Stringer, C. & Harris, K. D. Robustness of spike deconvolution for calcium imaging of neural spiking. bioRxiv 156786 (2017).

12. Wilt, B., Fitzgerald, J. E. & Schnitzer, M. J. Photon shot noise limits on optical detection of neuronal spikes and estimation of spike timing. Biophysical journal 104, 51–62 (2013).

13. Vogelstein, J. T. et al. Fast nonnegative deconvolution for spike train inference from population calcium imaging. Journal of neurophysiology 104, 3691–704 (2010).

14. Yaksi, E. & Friedrich, R. W. Reconstruction of firing rate changes across neuronal populations by temporally deconvolved ca 2+ imaging. Nature methods 3, 377 (2006).

15. Greenberg, D. S., Houweling, A. R. & Kerr, J. N. Population imaging of ongoing neuronal activity in the visual cortex of awake rats. Nature neuroscience 11, 749 (2008).

16. Sasaki, T., Takahashi, N., Matsuki, N. & Ikegaya, Y. Fast and accurate detection of action potentials from somatic calcium fluctuations. Journal of neurophysiology 100, 1668–1676 (2008).

17. Russakovsky, O. et al. Imagenet large scale visual recognition challenge. International Journal of Computer Vision 115, 211–252 (2015).

18. Adam-Bourdarios, C. et al. The higgs boson machine learning challenge. In NIPS 2014 Workshop on High-energy Physics and Machine Learning, 19–55 (2015).

19. Svoboda, K. & Project, G. Simultaneous imaging and loose-seal cell-attached electrical recordings from neurons expressing a variety of genetically encoded calcium indicators (2015).

20. Friedrich, J. & Paninski, L. Fast active set methods for online spike inference from calcium imaging. In Adv Neural Inf Process Syst 29, 1984–1992 (2016).

21. He, K., Zhang, X., Ren, S. & Sun, J. Deep residual learning for image recognition. In Proceedings of the IEEE conference on computer vision and pattern recognition, 770–778 (2016).

22. Szegedy, C. et al. Going deeper with convolutions (Cvpr, 2015).

23. Speiser, A. et al. Fast amortized inference of neural activity from calcium imaging data with variational autoencoders. In Advances in Neural Information Processing Systems, vol. (accepted) (2017).

24. Kuemmerer, M., Wallis, T. & Bethge, M. Information-theoretic model comparison unifies saliency metrics. Proceedings of the National Academy of Science 112, 16054–16059 (2015).

25. Kummerer, M., Wallis, T. S. A. & Bethge, M. Saliency benchmarking: Separating models, maps and metrics. arxiv (2017). URL https://arxiv.org/abs/1704.08615.

26. Reynolds, S., Schultz, S. R. & Dragotti, P. L. Cosmic: A consistent metric for spike inference from calcium imaging. bioRxiv 238592 (2017).

27. Freeman, J. Open source tools for large-scale neuroscience. Current Opinion in Neurobiology 32, 156–163 (2015).

28. Pachitariu, M. et al. Suite2p: beyond 10,000 neurons with standard two-photon microscopy. bioRxiv (2016). http://www.biorxiv.org/content/early/2016/06/30/061507.full.pdf.

29. Pedregosa, F. et al. Scikit-learn: Machine learning in Python. Journal of Machine Learning Research 12, 2825–2830 (2011).

30. Oord, A. v. d., Kalchbrenner, N. & Kavukcuoglu, K. Pixel recurrent neural networks. arXiv preprint arXiv:1601.06759 (2016).

31. Kingma, D. P. & Welling, M. Auto-encoding variational bayes. arXiv preprint arXiv:1312.6114 (2013).

32. Cho, K., Van Merriёnboer, B., Bahdanau, D. & Bengio, Y. On the properties of neural machine translation: Encoder-decoder approaches. arXiv preprint arXiv:1409.1259 (2014).

33. Glorot, X., Bordes, A. & Bengio, Y. Deep sparse rectifier neural networks. Aistats (2011).

34. He, K., Zhang, X., Ren, S. & Sun, J. Deep residual learning for image recognition. arXiv (2015). 1512.03385.

35. Abadi, M. et al. TensorFlow: Large-Scale machine learning on heterogeneous distributed systems. arXiv (2016). 1603.04467.

36. Graves, A. Generating sequences with recurrent neural networks. arXiv (2013). 1308.0850.

37. Otoro. Handwriting generation demo in TensorFlow. http://blog.otoro.net/2015/12/12/handwriting-generation-demo-in-tensorflow/ (2015). Accessed: 2017-5-18.

38. Chollet, F. et al. Keras. https://github.com/fchollet/keras (2015).

39. Barlow, R. E., Bartholomew, D. J., Bremner, J. & Brunk, H. D. Statistical inference under order restrictions: The theory and application of isotonic regression (Wiley New York, 1972).

40. Jewell, S. & Witten, D. Exact spike train inference via l0 optimization. arXiv:1703.08644 (2017).

41. van den Oord, A. et al. Wavenet: A generative model for raw audio. CoRR abs/1609.03499 (2016).

